# Generation of a PARPi-sensitive homozygous *BRCA1*-methylated OVCAR8 cell line using targeted CRISPR gene editing

**DOI:** 10.1101/2024.09.23.614644

**Authors:** Ksenija Nesic, Sally Beard, Lijun Xu, Olga Kondrashova, Cassandra J. Vandenberg, Alan F. Rubin, Fan Zhang, Alexander Dobrovic, Nicola Waddell, Clare L. Scott, Kristy Shield-Artin, Matthew J. Wakefield

## Abstract

Up to 17% of high grade serous ovarian carcinomas (HGSOC) harbour *BRCA1* promoter methylation (me*BRCA1*), making them susceptible to treatment with targeted PARP inhibitor (PARPi) therapy. Unfortunately, me*BRCA1* loss can be acquired following PARPi or platinum chemotherapy, resulting in *BRCA1* re-expression and PARPi resistance. Our understanding of me*BRCA1* stability in HGSOC is currently limited, in part due to a paucity of pre-clinical models with homozygous me*BRCA1*. Herein, we describe the generation of a several OVCAR8 cell line derivatives containing landing pad constructs, for future functional studies, and representing various *BRCA1* states, including a homozygous me*BRCA1* variant. Our PARPi resistant OVCAR8 has two methylated *BRCA1* copies and one unmethylated copy, enabling *BRCA1* expression. CRISPR-Cas9 gene editing was used to delete copies of the *BRCA1* gene in landing pad-containing clones of this cell line (A6 and H4). We produced one variant with deletion of all *BRCA1* copies (H4-53), and another with two copies deleted and only a single methylated gene copy remaining (A6-30 – validated further using nanopore long-read sequencing). These both lacked *BRCA1* gene expression and were sensitive to PARPi treatment. The A6-30 line was transplanted into immunocompromised mice to generate a xenograft model that retained homozygous me*BRCA1* and demonstrated some response to PARPi *in vivo*. Thus, using CRISPR gene editing we have created several novel isogenic HGSOC cell line models, including one with homozygous me*BRCA1*, that will support future studies of me*BRCA1* stability and PARPi resistance.

## Introduction

Poly (ADP-ribose) polymerase (PARP) inhibitors (PARPi) have revolutionized the treatment of ovarian and breast cancers, particularly in those with mutations in the *BRCA1* and *BRCA2* genes [1]. These genes play a critical role in homologous recombination (HR) DNA repair and, when defective, render cancer cells more susceptible to targeted PARPi-induced DNA damage. However, there are other HR defects beyond *BRCA1* and *BRCA2* mutations that can sensitize cancers to PARPi therapy. *BRCA1* promoter methylation (me*BRCA1*), a gene silencing mechanism that doesn’t involve gene mutation, is detected in a significant proportion of high grade serous ovarian carcinomas (HGSOC; ∼12% of cases [2, 3]) and triple negative breast cancers (TBNC; ∼15% of cases [4]). When homozygous, me*BRCA1* can functionally mimic a *BRCA1* mutation by silencing gene expression and, thus, impair HR DNA repair. The resulting HR deficiency (HRD) can sensitize cancer cells to PARPi therapy. However, loss of me*BRCA1* under treatment pressure has been observed in patient cancers and is associated with the development of PARPi resistance. This includes the development of heterozygous me*BRCA1*, as a single unmethylated allele of the gene is sufficient to restore HR and drive drug resistance [3].

Given a substantial number of breast and ovarian cancer patients have me*BRCA1* in their tumours, and me*BRCA1* states can significantly impact therapy responses, understanding how me*BRCA1* is regulated and maintained in these cancers is an important research question with clinical implications. Unfortunately, the field is faced with limited homozygously methylated cell line models available for study. Whether this is due to a survival pressure to maintain functional HR in culture is unclear. To our knowledge, WEHI-CS62 (derived by our group from a patient derived xenograft) is the only HGSOC cell line with confirmed homozygous methylation [3]. While the OVCAR8 cell line is known to have methylation of the *BRCA1* promoter, the zygosity of methylation appears to vary between cell line lineages. Originally isolated from a patient who had undergone multiple lines of therapy, the OVCAR8 cell line was characterised as cisplatin resistant [5]. OVCAR8 has since been described as having full gene silencing and PARPi responsive by some groups, while others have reported incomplete gene silencing in OVCAR8 associated with PARPi resistance [6–12]. Our group has studied a PARPi-resistant variant of OVCAR8 that has heterozygous me*BRCA1* – one of three gene copies has lost me*BRCA1*, the cell line has *BRCA1* gene expression and forms RAD51 DNA repair foci [3].

Using a series of OVCAR8 cell line variants with various *BRCA1* functional states, we have been able to confirm that our OVCAR8 variant is indeed resistant to PARPi due to a single unmethylated copy of *BRCA1*. Using CRISPR gene editing, we were able to create matched isogenic versions of OVCAR8 with either heterozygous me*BRCA1* (parental OVCAR8 cell line, 1/3 copies methylated), homozygous me*BRCA1* (two *BRCA1* gene copies deleted, single remaining copy is fully methylated) and *BRCA1* KO (full deletion of all *BRCA1* copies). We found that the homozygous me*BRCA1* and full *BRCA1* KO variants were sensitive to PARPi compared to the heterozygous me*BRCA1* parental line. We were also able to generate a cell line xenograft of the homozygous me*BRCA1* derivative, which showed some PARPi response. To maximise utility of this cell line, we incorporated a landing pad construct prior to CRISPR gene editing to facilitate downstream functional studies [13]. Our data indicate that versions of the OVCAR8 cell line with different *BRCA1* functional states and HR capacity may exist in the field. Other researchers should therefore carefully assess the *BRCA1* methylation status and gene expression of their own OVCAR8 cell lines and consider this when interpreting PARPi responses and HR repair status.

## Methods

### Ethics

All experiments involving animals were performed according to the Australian Code for the Care and Use of Animals for Scientific Purposes 8th Edition, 2013 (updated 2021), and were approved by the WEHI Animal Ethics Committee (2022.030).

### Cell lines and reagents

Rucaparib and the OVCAR8 cell line were kindly provided by Clovis Oncology, while niraparib tosylate (MK-4827 tosylate) was purchased from Assay Matrix. Our OVCAR8 cell line and all derivatives were grown in DMEM-based FV media as previously described [3]. HEK293T cells were obtained from ATCC and cultured in DMEM (Gibco, Cat #10569-010) with 10% FBS (Sigma-Aldrich, Cat #F9423). Cell lines were routinely tested and shown to be negative for Mycoplasma using the MycoAlert Mycoplasma Detection Kit (Lonza, Cat# LT07-118). Cells were passaged by detachment with 1 x Trypsin (Sigma-Aldrich, Cat #T4174). STR profiling [14] and WES (data not shown) were used to track and confirm cell line identity.

### Integration of landing pad construct

Lentiviral vectors were produced by transfection of 0.5×10^6^ HEK293T cells in a 6-well plate with 1.7 µg pLenti-Tet-coBxb1-2A-BFP_IRES-iCasp9-2A-Blast_rtTA3, 0.4 µg pRSV-Rev, 0.8 µg pMDLg/pRRE, 0.5 µg pMD2.G using CaCl_2_ and HEPES. Medium containing HEPES was replaced twice the following day, and supernatant collected 14 hours following transfection before passing through a 0.45 µm filter. OVCAR8 landing pad lines were generated by incubating 0.5×10^6^ OVCAR8 cells per well in a 12-well plate with viral supernatant diluted ¼ into culture media with 8 µg/ml polybrene (Merck, Cat #H9268) and spun for 2 hours, 32°C, 1000 x g. Media containing viral supernatant was removed six hours later and replaced with fresh media. Five days after transduction, 2 µg/ml doxycycline (Sigma-Aldrich, Cat #D9891) was added to the media. Eight days after transduction, cells were selected with 20 µg/ml blasticidine (Sigma-Aldrich, Cat #15205) and most cells were visually confirmed to have died off, supporting a MOI of less than one. After twelve days of selection, cells were assessed for BFP fluorescence, and individual BFP positive cells were sorted into single wells of a 96-well plate using a BD FACSAria Fusion (BD Biosciences). Landing pad lentivector pLenti-Tet-coBxb1-2A-BFP_IRES-iCasp9-2A-Blast_rtTA3 was a gift from Kenneth Matreyek (Addgene plasmid # 171588 ; http://n2t.net/addgene:171588 ; RRID:Addgene_171588). VSV-G envelope expressing plasmid pMD2.G was a gift from Didier Trono (Addgene plasmid # 12259 ; http://n2t.net/addgene:12259 ; RRID:Addgene_12259). Third generation lentiviral packaging plasmid pMDLg/pRRE was a gift from Didier Trono (Addgene plasmid # 12251 ; http://n2t.net/addgene:12251 ; RRID:Addgene_12251). Third generation lentiviral packaging plasmid pRSV-Rev was a gift from Didier Trono (Addgene plasmid # 12253 ; http://n2t.net/addgene:12253 ; RRID:Addgene_12253).

### Targeted CRISPR gene editing of *BRCA1*

crRNAs were designed using the Integrated DNA Technologies (IDT) “Design custom gRNA” tool to target downstream of the coding region and upstream of the CpG sites assessed in *BRCA1* promoter methylation analysis. *BRCA1* upstream guide sequence: AAAGAGCCAAGCGTCTCTCG. *BRCA1* downstream guide sequence: ACCGAGACTCATCAACTCAC. Both crRNAs were combined with Alt-R CRISPR-Cas9 tracrRNA, ATTO 500 (IDT, Cat #1075927) at equimolar amounts and incubated at 95°C for 5 min. RNP complex was formed by combining 1.2 µl of crRNA:tracrRNA duplex with 1.7 µl Alt-R S.p. HiFi Cas9 Nuclease V3 (IDT, Cat #1081061) and 2.1 µl of DPBS (Gibco, Cat #14190-144) and incubating at room temperature for 20 min. 1 × 10^5^ OVCAR8 landing pad cells were resuspended in 20 µl of nucleofector solution combined with supplement at 4.5:1 from the Amaxa SE Cell Line 4D-Nucleofector X Kit S (Lonza, Cat #V4XC-1032), before addition of 5 µl RNP complex and 1 µl of Electroporation enhancer. 25 µl of this mix was transferred to a nucleocuvette strip and run on program CA137 in a 4D Nucleofector (Amaxa). Cells were incubated in media overnight before sorting individual ATTO 550 positive cells (excitation 554nm, emission 575nm) using a BD FACSAria Fusion into 96-well cell culture plates.

### *BRCA1* deletion PCR screening

As individual *BRCA1* dual CRISPR clones approached confluence, cells were detached with trypsin, ¼ of cell volume was added to 20 µl dilution buffer with 0.5 µl DNARelease additive from Phusion Human Specimen Direct PCR Kit (Thermo Scientific, Cat #F-150BID) before incubation at 98°C for 2min. 1 µl of this product was used as input for PCR with 5 µl SYBR Green PCR Master Mix (Applied Biosystems, Cat #4309155), 0.25 µl 2 µM primer mix and 3.75 µl water. To detect intact *BRCA1* the following primer sequences were used, forward: GCCCCATGTCAGGCAAAGGGTC, reverse: TCTGCTTGCTTCCCGCCTGC. To detect deleted *BRCA1* the following primer sequences were used, forward: TACGTCATCCGGGGGCAGAC, reverse: TCTGCTTGCTTCCCGCCTGC. Cycling was carried out on the Quantstudio 12K Flex System at 95°C for 10min, 30 cycles of 95°C for 15s and 60°C for 1min*, followed by 95°C for 15s, 60°C for 1min* and 75°C for 5s increasing 0.1 second then hold* for 5s until 85°C (*data recorded at indicated steps). Positive control for PCR was generated by combining 0.01pg of synthetic gBlock fragment encompassing the regions flanking the two crRNA guide sites (IDT) with 0.5ng/µl genomic DNA isolated from parent cell line. Samples were screened for the presence of melt curves corresponding to *BRCA1* deletion and WT PCR products. One clone with the deletion band but no WT band was deemed to be a homozygous *BRCA1* deletion (OVCAR8 H4-53 *BRCA1^-/-/-^*). Heterozygous clones with both deletion and WT bands detected on PCR were further screened for *BRCA1* promoter methylation status, where OVCAR8 A6-30 (*BRCA1*^me/-/-^) was identified.

### Nanopore long-read DNA sequencing and analysis

DNA was extracted from OVCAR8 A6-30 (*BRCA1*^me/-/-^) cell pellets using the QIAamp DNA Mini kit according to the manufacturer’s protocol. The concentration of the extracted DNA was measured using the Qubit dsDNA high-sensitivity assay with a Qubit 3.0 fluorometer (Life Technologies). DNA purity was assessed using Implen NanoPhotometer® NP80. DNA size distribution was measured using genomic DNA quality assay on Agilent Tapestation. The extracted DNA was then fragmented, and size selected using Covaris g-TUBE at 5500RPM for 60 s. DNA Library was constructed using the Ligation Sequencing Kit (SQK-LSK114) and were sequenced using R10.4.1 flowcells on a PromethION device.

Read processing: Nanopore read data in POD5 file format were basecalled using Dorado (v0.7.1) with the pre-trained basecalling model dna_r10.4.1_e8.2_400bps_sup@v4.3.0. Trimming was performed based on NanoQC[15] reports, removing 40 nucleotides from the start and 20 nucleotides from the end of each read using Chopper (v0.7.0). Trimmed reads were aligned to a concatenated reference genome, which included the human reference genome GRCh38 and the pLenti-Tet-coBxb1-2A-BFP_IRES-iCasp9-2A-Blast_rtTA3 lentivirus construct[16]. The alignment was performed with minimap2 (v2.26)[17]. Aligned files were sorted and indexed by SAMtools (v1.17)[18]. The mapped OVCAR8 A6-30 (*BRCA1*^me/-/-^) reads were haplo-tagged by whatshap (v1.7)[19] based on phased variants generated by Clair3 (v1.0.4)[20].

Breakpoint detection: Two structural variant callers were used to identify the breakpoints around the *BRCA1* region and the sites of viral genome integration. The initial detection of events and their coordinates was performed using Severus (v1.0)[21] in tumor-only mode with default settings. Nanomonsv (v0.7.1)[22], run with a prebuilt control panel[23], was subsequently used to validate these findings. IGV (v2.16.0)[24] was used to visualise breakpoints.

### *BRCA1* methylation testing

DNA was extracted from snap-frozen cell pellets or CDX tissue using the QIAmp DNA Mini Kit (Qiagen, Cat #51304) or AllPrep DNA/RNA Mini Kit (Qiagen, Cat #80204) per manufacturer’s instructions. *BRCA1* methylation-specific high resolution melt analysis (MS-HRM) was performed as previously described [3]. *BRCA1* methylation sequencing (meNGS) was performed as previously described [25]. New primers specific to the *BRCA1* promoter region hg38 chr17:43125457 – 43125336 were designed for this assay (based on primers from [26], with Illumina sequencing adaptors added):

Forward primer (5’-3’), Illumina sequencing adaptors as bolded text:

**TCGTCGGCAGCGTCAGATGTGTATAAGAGACAG**TtgTtgTttagcggtagTTTTttggt;

Reverse primer (5’-3’), Illumina sequencing adaptors as bolded text:

**GTCTCGTGGGCTCGGAGATGTGTATAAGAGACAG**aAcctAtcccccgtccaAAaa

MeNGS data was analysed using the MethAmplicons package, using a 0.01 read frequency cut-off [25]. All samples were found to have an acceptably high bisulfite conversion efficiency (>98%), which reduces false positive methylation calls.

### *BRCA1* reverse transcriptase quantitative PCR

High quality RNA was extracted from snap-frozen cell pellets using the RNeasy Mini Kit (Qiagen, Cat #74104) or AllPrep DNA/RNA Mini Kit (Qiagen, Cat #80204) according to the manufacturer’s protocol. Resulting RNA was quantitated using the DeNovix Microvolume Spectrophotometer (DeNovix, Cat #DS-11). SuperScript III Reverse Transcriptase (Invitrogen, Cat# 18080085) was used to convert RNA to cDNA according to the manufacturer’s protocol. A volume of 2 µl cDNA diluted to 5 ng/µL was added to 5 µL SYBR Green PCR Master Mix, 2.5 µL molecular-grade H_2_O and 0.5 µL of the relevant 2 µM primer mix (previously described in [27]). Plates were incubated on the Quantstudio 12K Flex System (Thermo Fisher Scientific) at 95°C for 10min, 40 cycles of 95°C for 15s and 60°C for 1min*, followed by 95°C for 15s, 60°C for 15s and 95°C for 15s* (*Data recorded at indicated steps). Data were analysed using Design and Analysis Software 2.6.0 with *HPRT1*, *ACTB*, *SDHA* and *GAPDH* as endogenous reference genes, relative to OVCAR8 to produce RQ values, which represent fold change from OVCAR8. These values were then graphed in Prism 10 (GraphPad), and the standard deviation calculated.

### RNA sequencing and analysis

RNA QC, library preparation and sequencing were performed at the Australian Genomics Research Facility (AGRF). Sample quality was assessed using the RNA ScreenTape System (Agilent); all samples had a RIN of 10. Libraries were generated using the Illumina TruSeq Stranded mRNA library preparation, and sequencing was performed using the NovaSeq 6000 platform to produce 100bp paired end reads. Reads were mapped to human GRCh38 Ensembl 110 annotation using HISAT2 (v2.2.1) [28, 29]. BAM files were coordinate sorted using Samtools (v1.9) [30] before aligned reads were mapped to exons and quantified using HTseq (v3.0.3) [31]. Filtering was carried out and TMM normalization was performed [32]. Differential expression was carried out after voom transforming the counts using limma (v3.54.2) [33] to compare PARPi sensitive samples to resistant samples. A heatmap was produced from the top 30 DE genes using the R function pheatmap (v1.0.12).

### *In vitro* PARPi responses measured on the IncuCyte live cell imaging platform

Cells were plated in duplicate wells on Greiner 384 well plates, at 500 cells/well. The day following seeding, cells were treated with indicated doses of niraparib and incubated in a Sartorius IncuCyte S3 Live-Cell Analysis System where wells were imaged, using the 10x objective, once every 12 hours for up to 10 days. For growth analysis, a cell confluence mask was applied to all wells at each timepoint and plotted using the built-in IncuCyte software.

### Xenograft generation and *in vivo* treatments

The OVCAR8 A6-30 (*BRCA1*^me/-/-^) cell line xenograft model was established by resuspending 10 million cells in 1ml of a 50:50 mixture of Matrigel and DPBS. 100 µl was then injected subcutaneously per NOD.Cg*-Prkdc^scid^ Il2rg^tm1Wjl^/*SzJ (NSG; colony derived from Jackson Labs in 2018) recipient mouse (T1, passage 1) and mice were monitored for tumor formation. Tumors were harvested once tumor volume reached 700 mm^3^ and transplanted to new mice (T2) for treatments. Recipient mice bearing T2 (passage 2) tumors were randomly assigned to treatments when tumor volume reached 180-300 mm^3^. The regimen for rucaparib treatment was oral gavage once daily Monday-Friday for 3 weeks at 450 mg/kg for all models. Vehicle treatment was 0.5% methylcellulose oral gavage once daily Monday-Friday for 4 weeks. Tumors were harvested once tumor volume reached 700 mm^3^ or when mice reached ethical or end of experiment (120 days post treatment) endpoints. Data was plotted using the SurvivalVolume package [34].

## Results

### Generation of *BRCA1* deficient OVCAR8 variants

Experiments were conducted in OVCAR8 cell line derivatives containing a landing pad construct (Supplementary Figure 1) [13, 35] created for future gene integration/overexpression experiments. This Bxb1 recombinase-based landing pad with Tet inducible expression allows for HR-independent stable integration for reproducible expression and comparison between different constructs. OVCAR8 has three copies of the *BRCA1* gene. Targeted CRISPR gene editing was used to generate new *BRCA1*-deficient variants of the OVCAR8 cell line. Our OVCAR8 cell line, including derivatives harbouring the landing pad system, has *BRCA1* gene expression and is PARPi resistant due to heterozygous *BRCA1* methylation, where one gene copy is unmethylated: *BRCA1*^me/me/+^ (Figure 1A). Dual CRISPR guides targeting either end of the *BRCA1* (encompassing all exons and the gene promoter region) were used to create whole gene deletions (Figure 1B). Single cell sorting was used to generate various clones with different *BRCA1* deletion and methylation patterns.

**Figure 1.**
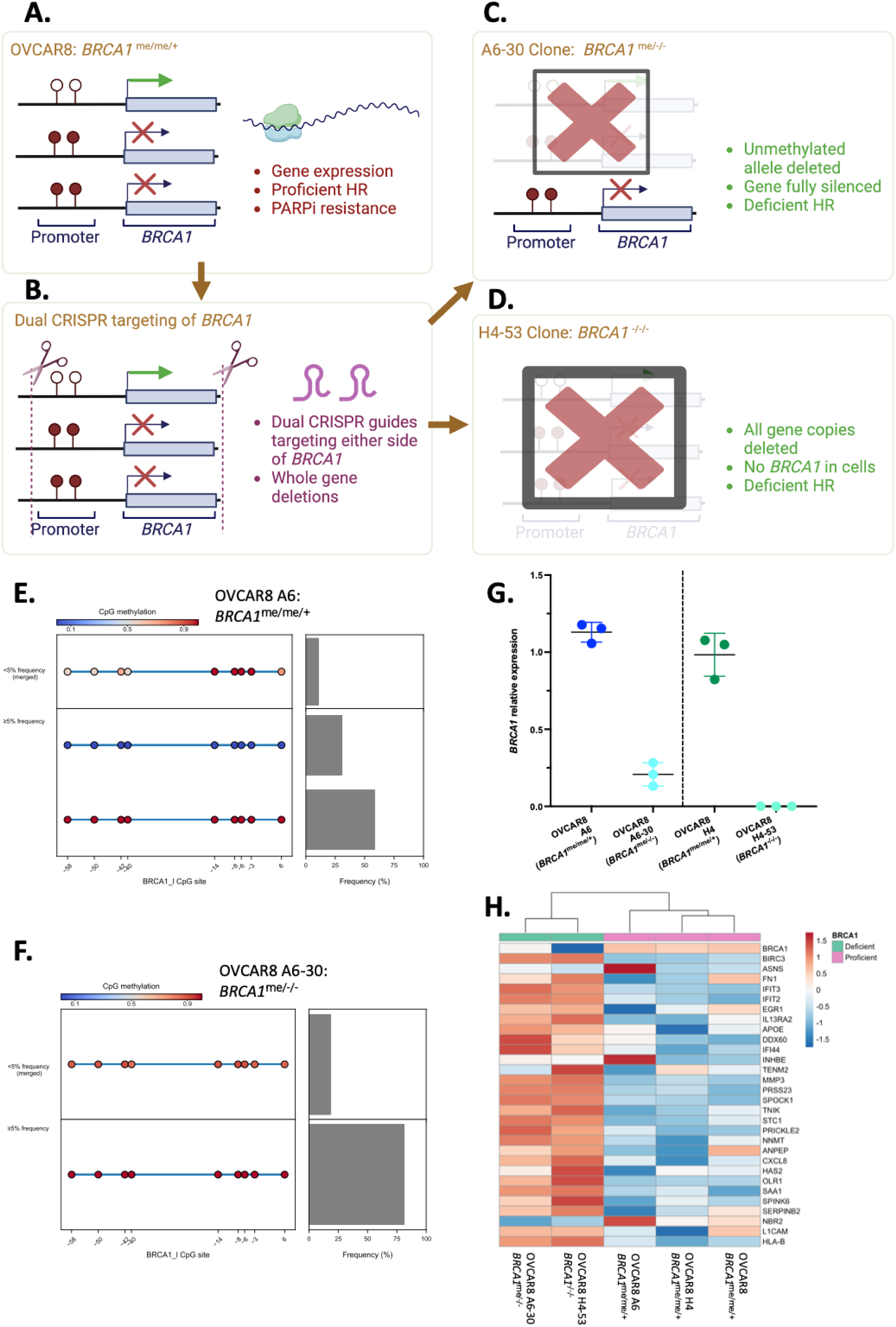
Generation of OVCAR8 landing pad derivatives with unique *BRCA1* deficiency states. (A) The OVCAR8 cell line in our laboratory, including A6 and H4 clones which contain a landing pad, has heterozygous *BRCA1* methylation (*BRCA1*^me/me/+^), with one of three gene copies lacking promoter methylation, permitting *BRCA1* gene expression and HR repair (as described in [3]). (B) We designed CRISPR guides to target either end of the *BRCA1* gene, including all exons and the gene promoter, and performed CRISPR gene editing in landing-pad containing OVCAR8 derivatives to generate clones with various *BRCA1* allele deletions and *BRCA1* deficiency states. (C) One of these resulting clones (A6-30, *BRCA1*^me/-/-^) had two copies of the gene deleted, leaving one copy with *BRCA1* promoter methylation, and was predicted to lack *BRCA1* gene expression, causing HR deficiency. (D) Another clone (H4-53, *BRCA1*^-/-/-^) was found to have deletion of all *BRCA1* copies, and was also predicted to lack *BRCA1* gene expression, driving HR deficiency. (E) Targeted *BRCA1* bisulfite sequencing confirmed heterozygous *BRCA1*^me/me/+^ methylation in the OVCAR8 A6 cell line (OVCAR8 with addition of landing pad construct), and (F) homozygous *BRCA1*^me/-/-^ methylation in the OVCAR8 A6-30 derivative following CRISPR. Circles represent CpG sites across the *BRCA1* promoter in regions known to be associated with silencing [3, 26], while values on x-axis indicate distance from the gene transcriptional start site. Colour scale represents 100% CpG methylation (red/1) to 0% methylation (blue/0). Epialleles present at a <5% frequency are all merged to show average CpG methylation (<5% merged), while those detected at a ≥5% frequency are presented individually. (G) The *BRCA1*^me/-/-^ A6-30 and *BRCA1*^-/-/-^ H4-53 derivatives were confirmed to have reduced or absent *BRCA1* mRNA transcript expression, respectively, using RT-qPCR (mean +/- standard deviation). Expression relative to average of four housekeeping genes. (H) RT-qPCR findings were supported by bulk RNA sequencing of the original OVCAR8 (*BRCA1*^me/me/+^), the A6 (*BRCA1*^me/me/+^) and H4 (*BRCA1*^me/me/+^) landing pad derivatives, and the *BRCA1*-deficient derivatives A6-30 (*BRCA1*^me/-/-^) and H4-53 (*BRCA1*^- /-/-^). Top differentially expressed genes between *BRCA1* proficient lines (OVCAR8, landing pad clone A6 and H4) compared to *BRCA1* deficient lines (A6-30 and H4-53) are presented on the right-hand side of the heat map of logCPM Z-scores scaled by rows.

Using this approach, one clone with homozygous (∼100%) *BRCA1* promoter methylation (Figure 1C – A6-30 clone, one methylated gene copy retained, *BRCA1*^me/-/-^) and another clone with loss of all *BRCA1* copies (Figure 1D, H4-53 clone, *BRCA1*^-/-/-^) were identified. The homozygous *BRCA1*-methylated *BRCA1*^me/-/-^ clone was derived from landing pad clone A6, and was thus named “OVCAR8 A6-30”. The *BRCA1*^-/-/-^ clone was derived from landing pad clone H4, and was thus named “OVCAR8 H4-53”. While not ideal that these lines were from different landing pad clones, the difficulty in growing *BRCA1* deficient cell lines left us with few options. These *BRCA1* deficient derivatives were identified using PCR-based screening, with results for A6-30 validated using long-read nanopore sequencing (Supplementary Figure 2), targeted *BRCA1* bisulfite sequencing (meNGS) and methylation-specific high resolution melt analysis (Figure 1E-F; Supplementary Figure 3). Nanopore sequencing confirmed the *BRCA1* methylation and copy number status of the OVCAR8 A6-30 *BRCA1*^me/-/-^ derivative, and also allowed identification of precise deletion breakpoints (Supplementary Figure 2). The effects of these unique *BRCA1* states on *BRCA1* mRNA gene expression were interrogated using reverse-transcriptase quantitative PCR (RT-qPCR). Compared to their respective parental landing pad lines, both *BRCA1*^me/-/-^ (A6-30) and *BRCA1*^-/-/-^ (H4-53) had little to no *BRCA1* gene expression (Figure 1G). Interestingly, *BRCA1*^me/-/-^ A6-30 appeared to have a very small amount of *BRCA1* gene expression, while the *BRCA1*^-/-/-^ H4-53 line had no detectable *BRCA1* mRNA transcripts. This was also reflected in bulk RNA sequencing of these cell lines (Figure 1H). The top 30 differentially expressed genes for OVCAR8 *BRCA1* deficient lines (H4-53 and A6-30) compared to parental lines (A6, H4 and OVCAR8) are presented (Figure 1H). While OVCAR8 A6 appears to be an outlier, it is not clear whether this is due to landing pad integration or features of the specific landing pad clone that formed this landing pad line.

### OVCAR8 variants with full *BRCA1* knock-out or *BRCA1* silencing are responsive to PARPi

With the new OVCAR8 *BRCA1*-deficient derivatives characterized, we interrogated the effects of each unique *BRCA1* deficiency on PARPi response. Parental landing pad cell lines OVCAR8 A6 and OVCAR8 H4 (both *BRCA1*^me/me/+^), and *BRCA1*-deficient derivatives OVCAR8 A6-30 (*BRCA1*^me/-/-^) and OVCAR8 H4-53 (*BRCA1*^-/-/-^), were treated with various doses of the PARPi niraparib and responses were measured every 12 hours over a 10-day time course on the IncuCyte live cell imaging platform (Figure 2A-D; Replicates in Supplementary Figure 4). Both *BRCA1*-deficient cell lines (*BRCA1*^me/-/-^ and *BRCA1*^-/-/-^) were clearly more sensitive to PARPi compared to the parental lines, with no cell growth observed at any tested PARPi niraparib doses in the *BRCA1*-deficient derivatives. It was observed that the OVCAR8 H4-53 (*BRCA1*^-/-/-^) and OVCAR8 A6-30 (*BRCA1*^me/-/-^) derivatives had slower growth compared to the corresponding *BRCA1*-expressing (*BRCA1*^me/me/+^) parental lines. This may be due to the importance of *BRCA1* for cell functions beyond HR DNA repair. The residual *BRCA1* mRNA detected in OVCAR8 A6-30 (*BRCA1*^me/-/-^), may permit better growth compared to OVCAR8 H4-53 (*BRCA1*^-/-/-^).

**Figure 2.**
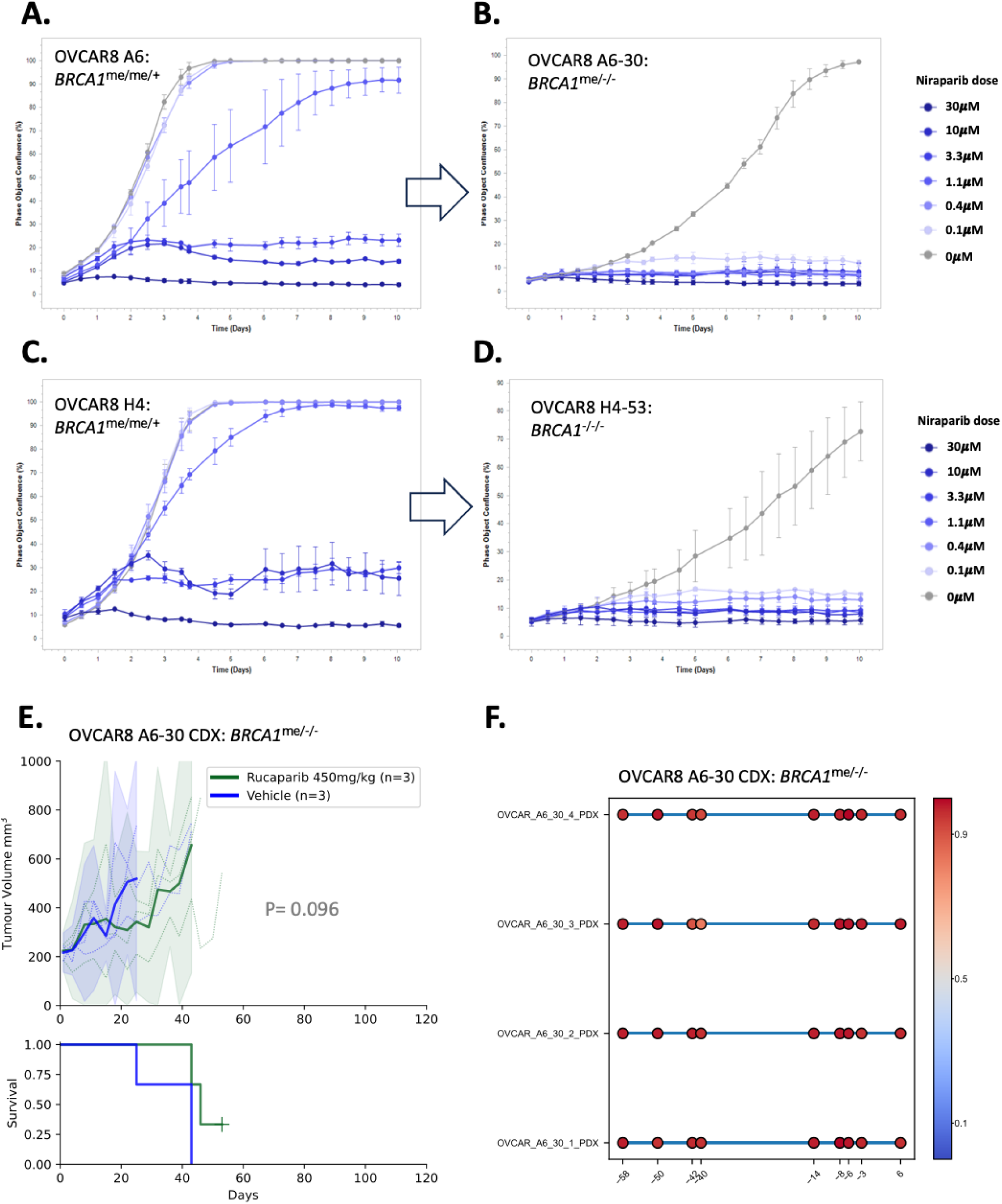
OVCAR8 derivatives with *BRCA1* deletion or silencing become PARPi responsive. Responses to the indicated doses of PARPi niraparib were assessed over 10 days (imaging every 12 hours) in (A) OVCAR8 A6 *BRCA1*^me/me/+^, (B) OVCAR8 A6-30 *BRCA1*^me/-/-^ with homozygous *BRCA1* methylation, (C) OVCAR8 H4 *BRCA1*^me/me/+^, and (D) OVCAR8 H4-53 *BRCA1*^-/-/-^, with full deletion of all *BRCA1* alleles. Data were from a representative experiment (mean and SEM of 3 technical replicates), with 2 additional experimental replicates presented in supplementary figure 4. (E) A cell line xenograft of OVCAR8 A6-30 (*BRCA1*^me/-/-^) was found to have minimal response to 450mg/kg PARPi rucaparib treatment *in vivo* compared to vehicle treatment (P= 0.096), despite (F) retaining homozygous methylation of *BRCA1* as assessed by targeted *BRCA1* bisulfite sequencing. Circles represent CpG sites across the *BRCA1* promoter in regions known to be associated with silencing [3, 26], while values on x-axis indicate distance from the gene transcriptional start site. Colour scale represents 100% CpG methylation (red/1) to 0% methylation (blue/0).

We explored whether these *BRCA1* deficient derivatives could form xenografts in immunocompromised NSG mice, and whether these xenografts are PARPi responsive. We transplanted the homozygous methylated derivative OVCAR8 A6-30 (*BRCA1*^me/-/-^) into NSG mice, then treated the cell line derived xenograft (CDX) with the PARPi rucaparib at a 450mg/kg dose. The xenografted cells readily grew in NSG mice, with slower growth in the PARPi rucaparib treated mice (Figure 2E); however, the response was not as impressive as the niraparib responses observed in the cell lines *in vitro* (Figure 2A-E; P= 0.096), with only a small shift in time to harvest between rucaparib-treated and vehicle-treated mice (46 vs 43 days, respectively). Whether this is a difference between the two PARPi, or a feature of the xenografted cells, needs to be explored further. The homozygous *BRCA1* methylation (*BRCA1*^me/-/-^) status of the PDX, in both PARPi treated and untreated samples, was confirmed using meNGS (Figure 2F; Supplementary Figure 5), suggesting that some other mechanism might be contributing to this difference in PARPi responses.

## Discussion

In this study, we successfully generated a series of unique OVCAR8 derivatives with unique *BRCA1* deficiency states (heterozygous methylation, fully methylated or fully deleted), that also contain landing pad loci for future functional genomics work [13, 35]. We confirmed that *BRCA1* expression is impaired in both the homozygously methylated and fully deleted derivatives, and that this loss of *BRCA1* leads to PARPi sensitivity in both cell lines. Given the paucity of *BRCA1* methylated HGSOC cell lines, this series of OVCAR8 with different HR DNA repair states will be invaluable for mechanistic studies of me*BRCA1* stability in HGSOC. Indeed, having both the derivative with *BRCA1* silencing and the other with full *BRCA1* deletion will be useful for studies of PARPi sensitivity and resistance mechanisms in these two distinct HR contexts. For example, interrogating which PARPi resistance mechanisms are unique to me*BRCA1* compared to full *BRCA1* deletion. We can now apply this approach to other cell lines that have heterozygous *BRCA1* methylation, or cell lines with other resistance mechanisms that only affect a single allele, such as secondary reversion or splice site mutations in HR genes [27].

Researchers also have the potential to use this set of cell lines to explore the incomplete dominance or haploinsufficiency of a single expressed wild-type *BRCA1* allele, where HR repair might be partially impaired or intermediate [36–38]. Indeed, this phenomenon might explain why different groups report different PARPi responses for the OVCAR8 cell line [3, 6–12]. This could be done by de-methylating the two silenced copies of *BRCA1* in the OVCAR8 cell line using DNA de-methylating agents, like decitabine or 5-azacytidine, or CRISPR-based approaches [39].

There are still aspects of these lines that will, however, require further interrogation. For example, why does the OVCAR8 A6-30 cell line have a low level of *BRCA1* expression, yet is supremely sensitive to PARPi *in vitro*? Also, why was the CDX of OVCAR8 A6-30 not as responsive to PARPi rucaparib as the corresponding cell line was to PARPi niraparib? While the CDX still retained homozygously methylated *BRCA1*, it is possible that the *BRCA1* gene expression might have been different between the two models. It may also be due to a difference in activity between niraparib and rucaparib. Niraparib generally has a higher selectivity for PARP1 and PARP2 compared to rucaparib [40], which might contribute to the difference we have observed. Given promising emerging data on the new generation of highly potent PARP1-specific inhibitors (*e.g.* AZD5305 [41, 42]), these inhibitors are also worth investigating in our cell lines and CDX.

The potential of the landing pad system in these cell lines warrants further exploration. By applying a well-established landing pad technology, we will be able to introduce and overexpress certain genes or gene variants in these cell lines using a straightforward recombinase strategy, enabling further characterization of HR pathway interactions and PARPi responses in these different *BRCA1* deficient contexts.

In summary, we have presented a CRISPR-based method of generating isogenic cell lines with distinct HR repair states by deleting specific copies of a given HR gene – in our case, targeting the *BRCA1* gene in the OVCAR8 cell line. The matched cell line derivatives we have generated in this study will be of great use for downstream studies of PARPi sensitivity and resistance mechanisms, particularly by also leveraging the presence of the landing pad locus in each cell line.

## Acknowledgments

We thank Silvia Stoev, Kathy Barber, Chloe Neagle, Steph Bound and Dan Fayle for technical assistance. This work was made possible through the Australian Cancer Research Foundation, the Victorian State Government Operational Infrastructure Support and Australian Government NHMRC IRIISS.

