## Supplementary Figure for "Generation of a PARPi-sensitive homozygous *BRCA1*-methylated OVCAR8 cell line using targeted CRISPR gene editing"

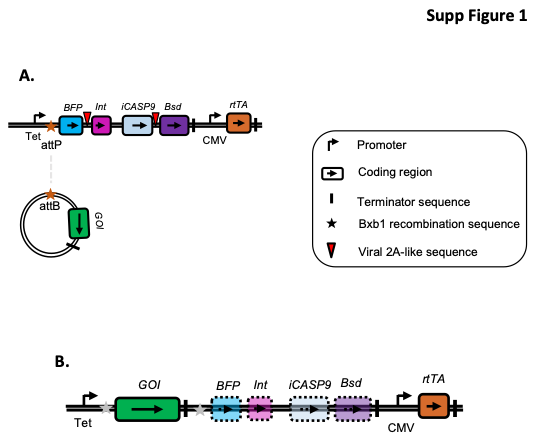

**Supplementary Figure 1. Structure of the landing pad safe harbour construct. (A)** Landing pad construct that is integrated into the genome of OVCAR8 A6 and H4. Tet is the tet inducible promoter. BFP is blue fluorescent protein. Int is Bxb1 integrase that catalyses recombination between attP site in landing pad and attB site in plasmid. IRES is internal ribosome entry site. iCASP9 is inducible caspase 9, which allows for negative selection with rimiducid. Bsd is blastidicin S deaminase. CMV is Cytomegalovirus promoter. rtTA is reverse tet transactivator. attP and attB are Bxb1 recombination sites. Viral 2A-like sequences allow co-translational separation of the polyprotein. **(B)** Genomic landing pad locus after integration of attB plasmid containing gene of interest. Tet inducible promoter now promotes inducible expression of the gene of interest (GOI). BFP, Int, iCASP9 and Bsd are switched off.

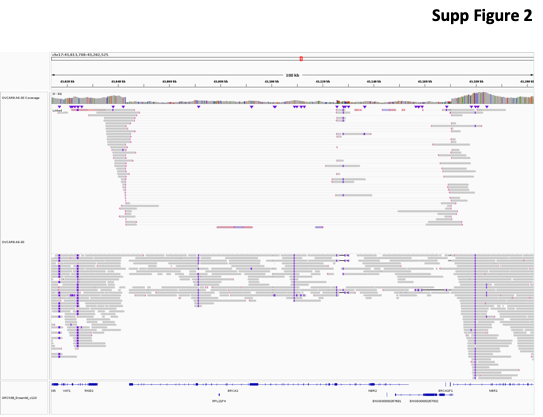

**Supplementary Figure 2. Nanopore long-read sequencing of OVCAR8 A6-30 derivative.**

IGV snapshot of the OVCAR8 A6-30 alignment shows two deletions within the *BRCA1* region: Chr17: 43042951-43170371 (reference read count = 7, variant read count =21) and Chr17: 43042952-43125519 (reference read count = 4, variant read count =13). In addition, a templated insertion cycle was detected by severus at chr17:43042970-43170340. Templated insertions are small DNA insertions that have been previously associated with HRD [1].

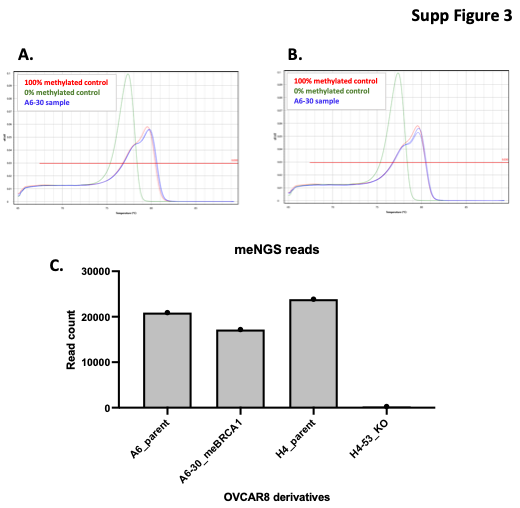

**Supplementary Figure 3. Methylation analysis of OVCAR8 derivatives. (A)** Methylation-specific high resolution melt was used to confirm homozygous *BRCA1* methylation in two aliquots of the OVCAR8-A6-30 cell line derivative. **(B)** Read counts for the targeted *BRCA1* bisulfite sequencing of derivative lines also support the presence of full *BRCA1* deletion in OVCAR8-H4-53, where no sequencing reads were detected.

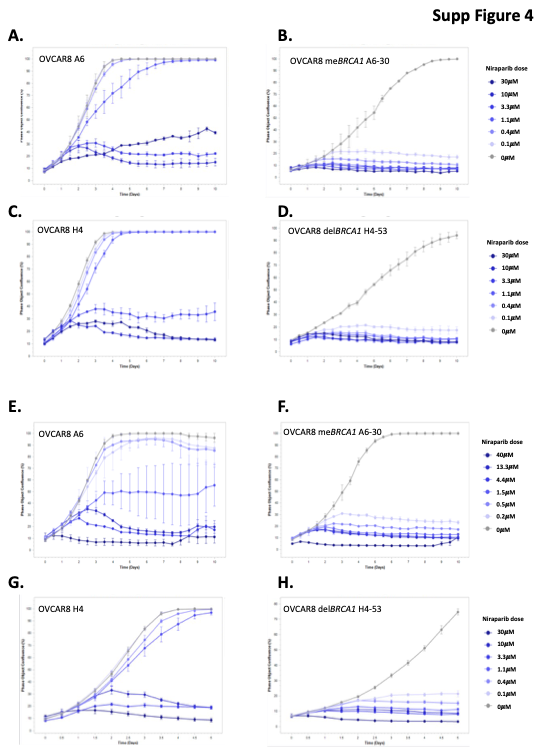

**Supplementary Figure 4. Replicate data of niraparib treatment experiments in OVCAR8 derivatives.** Responses to the indicated doses of PARPi niraparib were assessed over 10 days (imaging every 12 hours) for replicate 2 in **(A)** OVCAR8-A6, **(B)** OVCAR8-A6-30 with homozygous *BRCA1* methylation (me*BRCA1*), **(C)** OVCAR8-H4, and **(D)** OVCAR8-H4-53 with full deletion of all *BRCA1* alleles (del*BRCA1*). Replicate 3 for **(E)** OVCAR8-A6, **(F)** OVCAR8-A6-30 with homozygous *BRCA1* methylation (me*BRCA1*), **(G)** OVCAR8-H4, and **(H)** OVCAR8-H4-53 with full deletion of all *BRCA1* alleles (del*BRCA1*) used slightly different niraparib doses, but demonstrated the same trends.

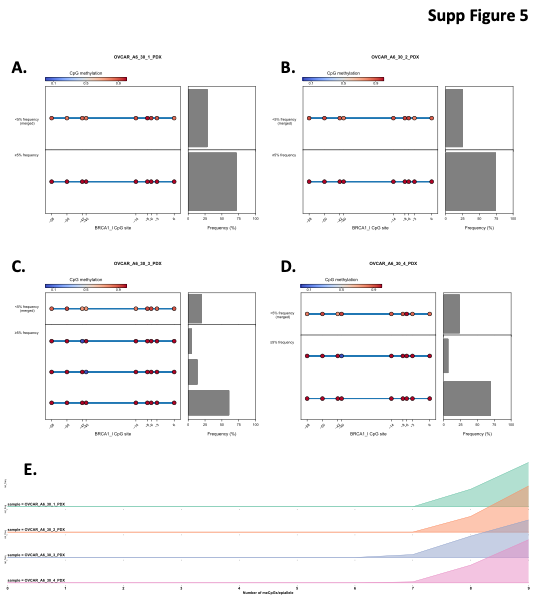

**Supplementary Figure 5. Homozygous *BRCA1* methylation confirmed in both rucaparib and vehicle treated OVCAR8-A6-30 xenograft samples.** Targeted *BRCA1* bisulfite sequencing confirmed homozygous *BRCA1* methylation in the OVCAR-A6-30 cell line xenograft in either **(A-B)** vehicle-treated and **(C-D)** rucaparib-treated tumour samples. Circles represent CpG sites across the *BRCA1* promoter in regions known to be associated with silencing [2, 3], while values on x-axis indicate distance from the gene transcriptional start site. Colour scale represents 100% CpG methylation (red/1) to 0% methylation (blue/0). **(E)** The absence of unmethylated epialleles in samples from A-D was confirmed by plotting the frequency of epialleles with 0 methylated CpGs to epialleles with all 9 methylated CpGs (x-axis).
